# The influence of early life exposures on the infant gut virome

**DOI:** 10.1101/2023.03.05.531203

**Authors:** Yichang Zhang, Josué L. Castro-Mejía, Ling Deng, Shiraz A. Shah, Jonathan Thorsen, Cristina Leal Rodríguez, Leon E. Jessen, Moïra B. Dion, Bo Chawes, Klaus Bønnelykke, Søren J. Sørensen, Hans Bisgaard, Sylvain Moineau, Marie-Agnès Petit, Jakob Stokholm, Dennis S. Nielsen

## Abstract

Large cohort studies have contributed significantly to our understanding of the factors that influence the development of the bacterial component of the gut microbiome (GM) during the first years of life. However, the factors that shape the colonization by other important GM members such as the viral fraction remain more elusive. Most gut viruses are bacteriophages (phages), i.e., viruses attacking bacteria in a host specific manner, and to a lesser extent, but also widely present, eukaryotic viruses, including viruses attacking human cells. Here, we utilize the deeply phenotyped COPSAC2010 birth cohort consisting of 700 infants to investigate how social, pre-, peri- and postnatal factors may influence the gut virome composition at one year of age, where fecal virome data was available from 645 infants.

Among the different exposures studied, having older siblings and living in an urban vs. rural area had the strongest impact on gut virome composition. Differential abundance analysis from a total of 16,118 viral operational taxonomic units (vOTUs) (mainly phages, but also 6.1% eukaryotic viruses) identified 2,105 vOTUs varying with environmental exposures, of which 5.9% were eukaryotic viruses and the rest was phages. Bacterial hosts for these phages were mainly predicted to be within the *Bacteroidaceae, Prevotellaceae*, and *Ruminococcaceae* families, as determined by CRISPR spacer matches. Spearman correlation coefficients indicated strong co-abundance trends of vOTUs and their targeted bacterial host, which underlined the predicted phage-host connections. Further, our findings show that some gut viruses encode important metabolic functions and how the abundance of genes encoding these functions is influenced by environmental exposures. Genes that were significantly associated with early life exposures were found in a total of 42 vOTUs. 18 of these vOTUs had their life styles predicted, with 17 of them having a temperate lifestyle. These 42 vOTUs carried genes coding for enzymes involved in alanine, aspartate and glutamate metabolism, glycolysis-gluconeogenesis, as well as fatty acid biosynthesis. The latter implies that these phages could be involved in the utilization and degradation of major dietary components and affect infant health by influencing the metabolic capacity of their bacterial host.

Given the importance of the GM in early life for maturation of the immune system and maintenance of metabolic health, these findings provide a valuable source of information for understanding early life factors that predispose for autoimmune and metabolic disorders.

## Introduction

Early life gut microbiome (GM) establishment plays a fundamental role in shaping host physiology and health^1,2^ with early life GM imbalances being linked to onset and progression of chronic diseases later in life, such as obesity^3^, diabetes^4,5^, and asthma^2^.

To date, GM research has generally focused on understanding the importance of the bacterial GM component, but recent findings indicate that the vast and diverse population of viruses found in the gut (collectively called the “virome”) also play a prominent role in gut microbial ecology^6–9^. Amidst these biological entities, bacterial viruses, also termed bacteriophages (phages), are the most diverse and abundant particles of the GM^9–11^ and they represent a major reservoir of genetic diversity influencing not only GM composition, but also the GM metabolic potential^12,13^. Disease-specific alterations in the gut virome have been reported in several chronic conditions^14^ such as inflammatory bowel disease^15^, colorectal cancer^16^, necrotizing enterocolitis in preterm infants^17^, severe acute malnutrition^18^, type-1 diabetes^19,20^ and other autoimmune diseases such as rheumatoid arthritis^21^. The role of the gut virome in shaping the GM is underlined by the observation that fecal virome transfer from healthy donors to recipients with a dysbiotic GM prevent or ameliorate symptoms associated with metabolic^22^ and gastrointestinal^23,24^ disorders.

While various early-life factors such as birth mode, siblings, diet and exposure to antibiotics has been found to influence development of the gut bacterial populations^1,25^, little is known about which factors shape the gut virome. The few attempts that have characterized the gut virome early in life have revealed that its composition is highly dynamic^26–28^, affected by delivery mode^6^ and the first bacterial colonizers^29^ as well as being enriched in phages belonging to the *Microviridae* family^10,27^. Moreover, its transmission-dynamics after birth follows a stepwise assembly, with breastfeeding playing a protective role against eukaryotic viral infections^30,31^. Understanding how environmental exposures and phenotypes intertwine the vector space conformed by viruses, bacteria, host, and their functional attributes remains an unsolved task.

In a recent detailed investigation of the infant gut virome, we showed a massive diversity of hitherto undescribed phages^9^. In this cross-sectional study of the gut virome of 645 infants at one year of age enrolled in the COPSAC2010 cohort^32^ more than ten thousand viral species distributed over 248 viral families and 17 viral order-level clades were detected. Here we investigate how social, pre-, peri- and postnatal factors influence the gut virome composition at one year of age. Our findings demonstrate how early life exposures are linked to the abundance of specific viruses, as well as their co-abundance and concordance with their predicted bacterial hosts. Metabolic functions encoded in the genomes of these viruses displayed enrichment of genes important for bacterial physiology in response to exposures, some of which are likely associated with dietary elements (e.g., degradation of complex carbohydrates) and others that may influence infant growth and health.

## Results

### Composition of DNA viruses in the gut of Danish infants

A total of 645 stool samples from 1-year old infants in the COPSAC2010 cohort^32^ were obtained and analyzed^9^. Virions were isolated, concentrated and their genome was sequenced using a shotgun metagenome strategy^9,33^. Following assembly, a total of 16,118 species-level clustered viral representative contigs (here termed viral Operational Taxonomic Units – vOTUs) were obtained. Around 70% of the vOTUs were affiliated to five viral classes (*Arfiviricetes, Caudoviricetes, Faserviricetes, Malgrandaviricetes* and *Tectiliviricetes*) (Figure 1A and 1I). Almost 18.8% of the vOTUs (n=3,029) were considered putative satellite phages as contigs lacked genes coding for structural proteins but encoded other viral proteins (e.g., integrases or replicases) and were conserved in size and gene content across multiple samples. In addition, 11.8% of the vOTUs (n=1,895) were categorized as unclassified viral fragments (Figure 1A).

**Figure 1.**
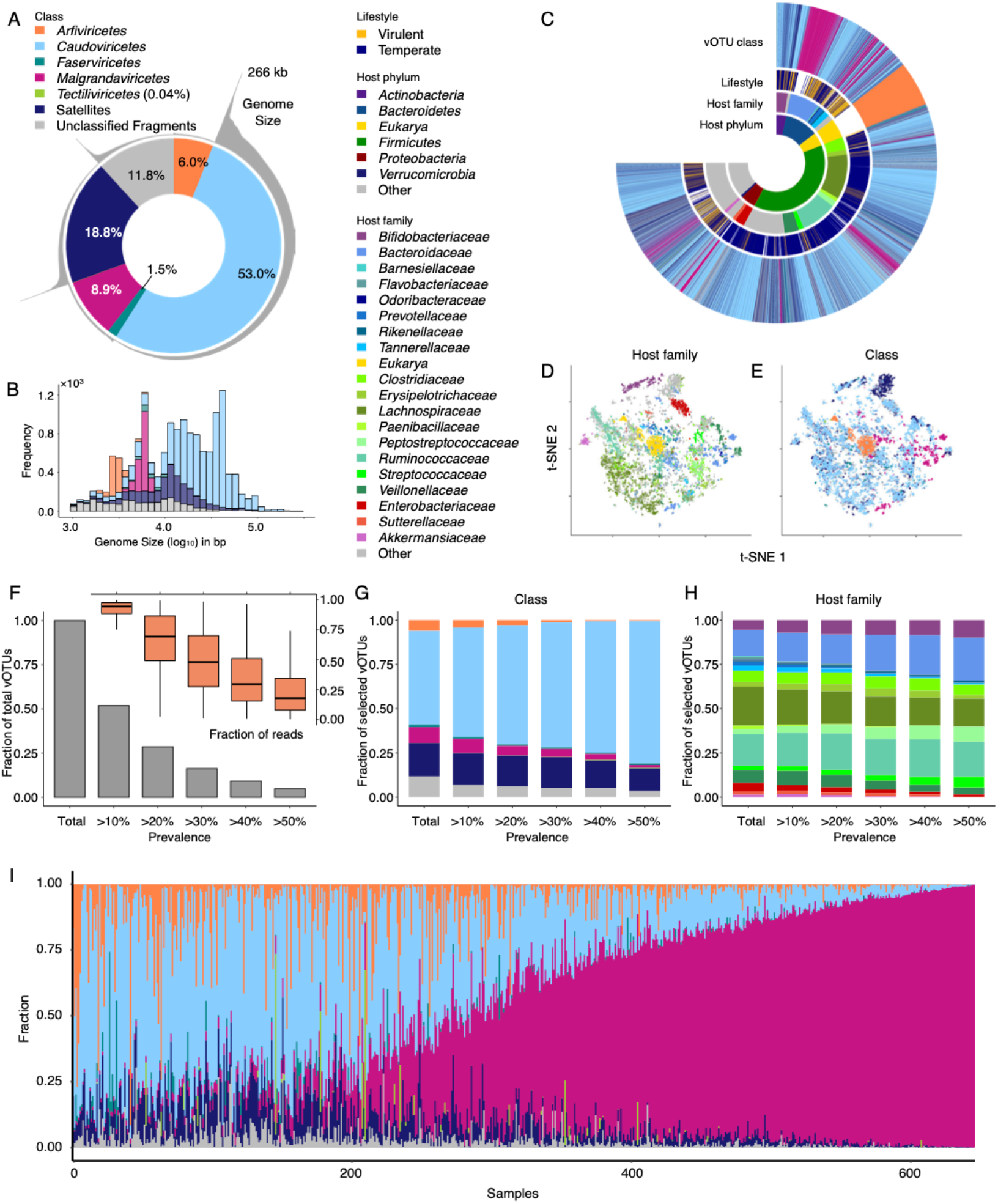
Virome structure of the infants enrolled in the COPSAC2010 cohort. A) Distribution of the 16118 vOTUs identified colored by their taxonomic class annotation. B) Cumulative frequency of viral genomes (kb) identified by their taxonomic class annotation. C) Circular diagram showing the distribution of vOTUs colored by their targeted bacterial hosts (at phylum and family levels), viral class and lifestyle. D-E) t-Stochastic Neighbor Embedding (t-SNE) plots clustering *tetra-mer* vOTUs profiles identified according to host family (D) and viral class (E). F-H) Percentage of vOTUs that appear at a specific prevalence (F), and vOTUs’ distribution colored by their taxonomic class (G) and host family (H). I) Relative abundance of vOTUs across all samples at the class level. Samples were sorted by *Malgrandaviricetes* abundance.

The largest genomes (>10 kb) were observed among *Caudoviricetes*, which constituted the vast majority of vOTUs (Figure 1B). The genomes, dominated by *Caudoviricetes* (tailed, double-stranded DNA phages) and *Malgrandaviricetes* (non-tailed, single-stranded DNA phages), followed a bi-/multi-modal distribution (Hartigans’ Dip test, P < 0.0001) based on their genome sizes (Figure 1B).

Bacterial hosts as well as lifestyle (temperate/virulent) of the vOTUs were predicted using CRISPR spacers and the presence of integrases^9^, respectively (Figure 1C). Because phages tend to have comparable *k*-mer frequencies to those of their hosts^34,35^, we also performed dimensionality reduction on tetramer vectors to confirm global host associations as a complement to our viral taxonomy^9^. Using unsupervised stochastic neighbor embedding (t-SNE) dimensionality reduction, vOTUs targeting the same hosts as determined by CRISPR spacers (Figure 1D) or belonging to the same viral classes (Figure 1E) were found to clearly cluster together. Previously only *Enterobacteriaceae* and *Bacteroidetes* have been shown to be the hosts of non-tailed *Malgrandaviricetes^36^*, but when examining the bacterial hosts, we observed that in addition to *Bacteroidetes*, also *Ruminococcaceae, Clostridiaceae, Erysipelotrichaceae* and *Sutterellaceae* are predicted as hosts of *Malgrandaviridetes* viruses (Figure 1C and S1). With respect to lifestyle, *Streptococcaceae* and most families of the *Bacteroidetes* have a greater proportion of vOTUs recognized as virulent than temperate (Figure 1C and Table S2).

The distribution of vOTUs was very individual-specific, with less than 5% of vOTUs appearing in more than 50% of the samples (Figure 1F). However, this still adds up to around 800 vOTUs that are shared among a majority of infants and representing, on average, more than 20% of the reads (Figure 1F). The proportion of vOTUs classified as *Caudoviricetes* (Figure 1G) as well as those infecting *Bacteroidaceae* and *Bifidobacteriaceae* (Figure 1H) increased as a function of prevalance.

### Environmental exposures influence viral diversity

A range of pre-, peri-, and postnatal as well as social factors were recorded for the enrolled infants and their families (Supplementary Table S1). Having older siblings was associated with higher vOTU richness (linear mixed model, *P* = 0.048, estimate = 69.14, 95% CI = [0.58, 137.52]) and lower evenness (Shannon *H’*) (linear mixed model, *P* = 0.003, estimate = −0.30, 95% CI = [−0.50, −0.10]) (Figure 2A-B and 2E-F) at one year of age. Likewise, a higher birth weight was linked to higher vOTU richness (linear mixed model, *P* = 0.007, estimate = −85.76, 95% CI = [−153.98, −17.56]) (Figure 2A and 2E). Dietary factors were also found to influence the gut virome at one year of age, with late introduction of eggs in the diet being associated with lower viral evenness (Shannon *H’*) (linear mixed model, *P* = 0.012, estimate = 0.25, 95% CI = [0.05, 0.45]) (Figure 2A and 2F). The mothers were enrolled in a nested randomized placebo-controlled trial of fish oil to the mothers during the third trimester of pregnancy^37,38^. Receiving fish oil during pregnancy was associated with increased gut vOTU richness (linear mixed model, *P* = 0.038, estimate = 71.60, 95% CI = [3.90, 139.22]) of the infants at one year of age (Figure 2A). The design also examined the difference in vitamin D between high and standard doses^39^, which had no effect on the viral community in our analysis. Interestingly, other factors that have been found to influence the bacterial GM component during infancy such as birth mode, use of antibiotics, and duration of exclusive breastfeeding did not seem to influence gut virome alpha-diversity measures at one year of age in this cohort (Figure 2A-B).

**Figure 2.**
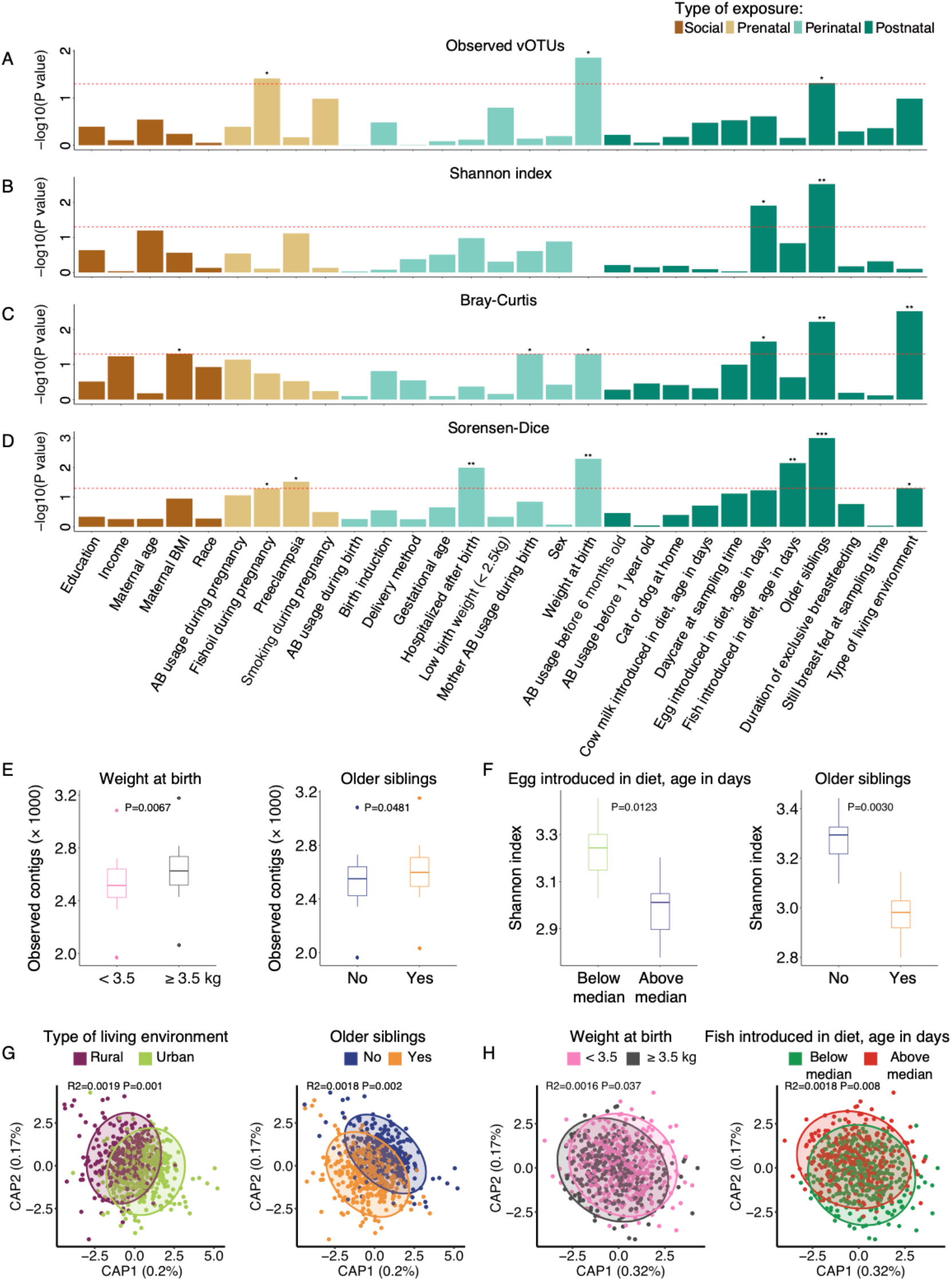
Virome diversity and composition covariates with early life exposures. A-D) Barplot showing the strength of associations (-log10 p-value) of the alpha diversity metrics Observed vOTUs (A) and Shannon Index (B) across different exposures (linear mixed model) as well as beta diversity using distance-based redundancy analysis (db-RDA) on Bray-Curtis dissimilarity (C) and Sorensen-Dice distance (D) matrices. E-F) Distribution of Observed vOTUs for weight at birth and siblings (E) and Shannon Index for siblings and dietary introduction of egg (F). Dietary introduction of egg is indicated in days. G-H) db-RDA constrained-components based on Bray-Curtis distances for location and siblings (G), and Sorensen-Dice distances for weight at birth and dietary introduction of fish (H).

Regarding virome composition, Bray-Curtis dissimilarity analysis (weighted measure, which is therefore mainly influenced by more abundant vOTUs) showed a link (PERMANOVA, *P* = 0.049, R2=0.0016) between maternal body mass index (BMI) and virome composition at one year of age (Figure 2C, 2G and S1A); while Sørensen-Dice distance (unweighted binary metric and therefore mainly influenced by more rare vOTUs) revealed that a number of pre- and perinatal exposures were linked with virome composition differences (PERMANOVA, *P* ≤ 0.05), namely weight at birth, fish oil supplementation during pregnancy, hospitalization after birth, and preeclampsia (Figure 2D, 2H and S2B). Both Bray-Curtis and Sørensen-Dice metrics showed significant differences in virome composition for children having older siblings (PERMANOVA, *P* = 0.006, R2=0.0018 and *P* = 0.001, R2=0.0029 for Bray-Curtis and Sorensen-Dice, respectively), and whether the family was living in an urban or a rural area (PERMANOVA, *P* = 0.003, R2=0.0019 and *P* = 0.049, R2=0.0016 for Bray-Curtis and Sorensen-Dice, respectively) (Figure 2C-D, 2H and S2A-B).

### Environmental exposure variables influence the abundance of specific vira

Subsequently, we determined how the distribution of vOTUs differed between the nine exposures (Figure 2C-D) found to significantly influence overall gut virome composition (preeclampsia was not included due to highly unbalanced sample size, see Supplementary Table S1). A total of 2,105 differentially abundant vOTUs affiliated to 173 viral families and 19 families of bacterial hosts were identified by DESeq2, with having older siblings being associated with 822 differential abundant vOTUs, while being hospitalized after birth being associated with 212 differential abundant vOTUs (Figure 3). For perinatal covariates, vOTUs differing in abundance were predicted to infect a range of different hosts, but interestingly revealed a pronounced lower abundance towards those infecting *Bacteroidaceae*, *Ruminococcaceae* and *Streptococcaceae* associated with with maternal antibiotic usage and hospitalization after birth (Figure 3). Postnatal factors like specific dietary patterns (late introduction of eggs in the diet), presence of older siblings in the house and living in a rural environment, were associated with a higher abundance of vOTUs infecting *Bifidobacteriaceae, Bacteroidaceae, Prevotellaceae, Tannerellaceae, Ruminococcaceae* and *Sutterellaceae*.

**Figure 3.**
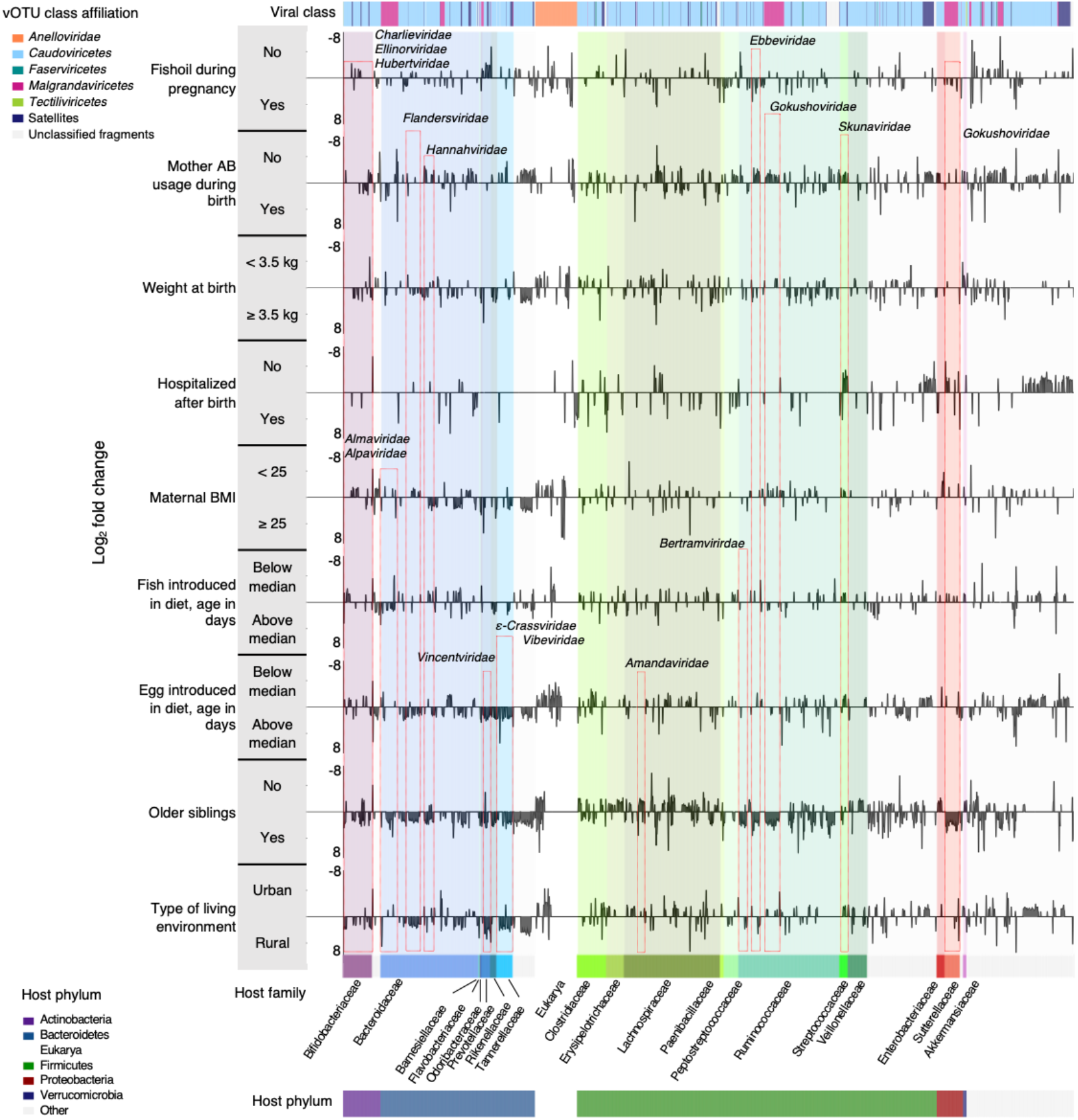
Viral host family, relative abundance and lifestyle associate with environmental exposures at one year of age. Visualization of differential abundance analysis of 2105 vOTUs across the nine exposures significantly associated with virome diversity and composition. Log_2_ fold change panel displays the change in abundance between the two groups for each exposure. The viral families to which vOTU belongs, surrounded by red boxes, are labeled. Adjusted P ≤ 0.001 and Log_2_-Fold changes ≥ |1| were used to select differentially abundant vOTUs.

To further integrate these findings in the context of the gut bacterial component, we used 16S rRNA gene (V4 region) amplicon sequencing (bacterial OTUs - bOTUs) data previously published for this cohort(Stokholm et al. 2018) to determine virus-host co-abundances. Spearman correlation coefficients (ρ) were calculated between the abundance of the above identified differentially different abundant vOTUs and bOTUs across samples. Only bOTUs that were strongly associated (ρ≥0.3) with at least one vOTU were retained. If a vOTU was correlated with a bOTU, the bOTU family tended to be consistent with the predicted host family of the vOTU (Figure S3A). These virus-host co-abundances indicate there is a high degree of inter-relatedness between phages and their host in response to environmental exposures. This was supported by the fact that the same perinatal and postnatal covariates were also significantly associated with bOTU diversity and composition (Figure S4A-D). Overall, among the 91 co-abundant vOTUs (ρ≥0.3), vOTUs that infect the same bacterial host family were in most cases closely related genetically, indicating a high degree of co-evolution between bacterial hosts and the phages that infect them (Figure S3B). To confirm the above-mentioned findings, we repeated the analysis of virus-host co-abundances using shotgun metagenomic data from the same cohort^40^. We found again that viruses and their bacterial hosts were highly correlated supporting the same conclusion as above (Figure S4E).

### Functional profiles of gut viruses are linked with environmental exposures

Differentially abundant vOTUs were subjected to gene (open reading frame, ORF) prediction, and annotated based on KEGG Orthology (KO) using KofamScan^41^. As seen from figure S5A, 0.82% of genes matched known metabolism-related orthologs, while the remaining genes with KO assignments (8.48% of predicted genes) encoded genes related to genetic information processing and signaling and cellular processes, representing typical viral-associated traits required to accomplish replication^42^. The remaining 90.7% of the predicted genes were not annotated by the database.

Next, we focused on determining genes with metabolic functions having the potential to enhance host fitness and drive metabolic reprogramming of the bacterial host^43^. The gut virome of infants with older siblings were enriched in genes related to O–antigen nucleotide sugar biosynthesis and seleno-compound metabolism, while infants without siblings were enriched in genes related to carbon fixation in photosynthetic organisms (Fisher’s exact test, P < 0.05; Figure 4A) (the link to photosynthetic microorganisms may be caused by the KEGG database not being optimized for vira). The gut of infants living in rural areas or that were introduced to eggs in their diet later in life (above the median age when eggs were introduced in the diet) were enriched in viral encoded genes associated with glycolysis/gluconeogenesis and O-antigen nucleotide sugar biosynthesis, whereas the gut of infants living in urban areas or that were introduced to eggs relatively early in life were enriched in viral genes associated with thiamine metabolism. Infants with birth weight above the median or whose mothers were not obese also encoded genes involved in diverse pathways involved in e.g. vitamin synthesis. Further, the gut virome of infants whose mothers received fish oil during pregnancy or were prescribed antibiotics during delivery encoded genes related to purine metabolism (Figure 4A).

**Figure 4.**
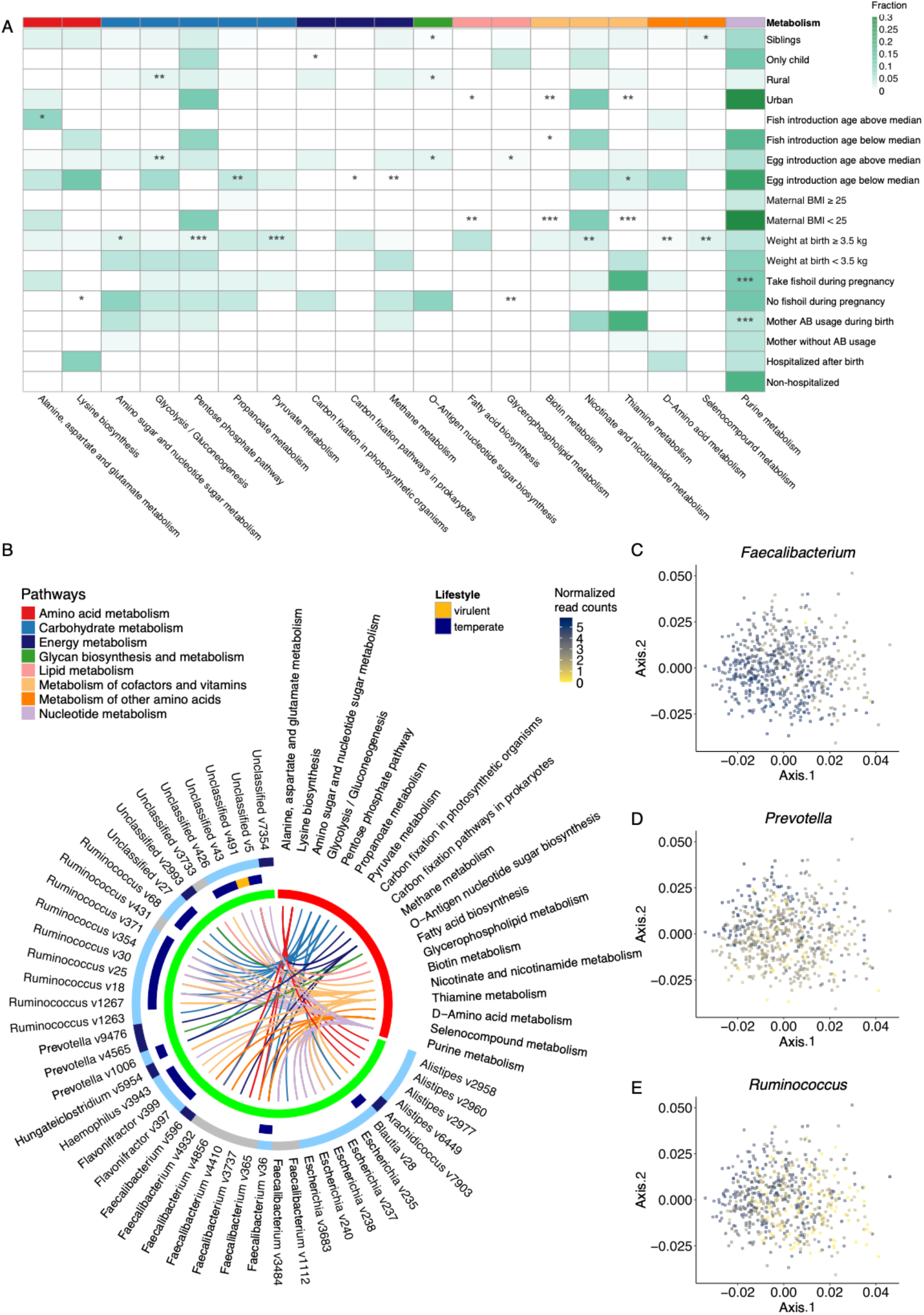
Abundance of phage accessory genes differ in connection with exposures. A) Abundance of genes (3^rd^ level KEGG pathway) in the virome of infants with significant (P ≤ 0.05) enzymatic enrichments that are associated with the presence of siblings and residential location. B) Viral host families that contribute to metabolism pathways. C-E) Bacterial points extracted from the procrustes analysis (E). The points are colored according to the abundance of the specific genus in each sample.

To determine how virally encoded gene functions associate with the microbial composition, we linked back enriched genes to the vOTU of origin (Figure 4B). 94% of lifestyle predicted vOTUs (n=17) were temperate. Genes associated with two classes of amino acid metabolisms (i.e. alanine, aspartate and glutamate metabolism and lysine biosynthesis) were conserved across *Alistipes* and *Faecalibacterium* targeting vOTUs, respectively. In addition, multiple carbohydrate metabolism enzyme encoding genes were found to be widely encoded by *Blautia*, *Prevotella*, *Ruminococcus* and *Faecalibacterium* targeting vOTUs. These encoded enzymes including L-lactate dehydrogenase, ribose-phosphate pyrophosphokinase and aldose 1–epimerase (Figure S5D). Energy metabolism genes were found in *Prevotella and Faecalibacterium* targeting vOTUs, while nicotinate and nicotinamide metabolism genes were mapped in *Ruminococcus* and *Escherichia* targeting vOTUs (Figure 4B).

Phage-host co-abundance (Figure S3A), was further confirmed by Procrustes analysis. The linking of the virome and bacteriome compositions revealed a strong correlation one to another (*P* < 0.001, *r* = 0.52) (Figure S5B-C). The cumulative abundance of all bOTUs belonging to the bacterial genera *Ruminococcus, Prevotella* and *Faecalibacterium*, which were found to be the main bacterial host of vOTUs carrying the above metabolic genes (Figure 4B) and having previously been reported to be highly associated with stable viral communities^44^, was highly correlated with rural vs. urban living and having older siblings (Figure 4C-E). These results emphasize the potential role of phage-host association in metabolic regulation.

## Discussion

The gut of healthy newborns is usually devoid of viruses at birth, but it is rapidly colonized afterwards^27,30^. Still relatively few studies have focused on the assembly of the gut virome within the first year of life and the factors that influence it^6,26,27,30^ and even less is known about the environmental exposures that shape the gut virome.

Here, we leveraged a massive gut virome dataset from healthy infants at 1-year of age, and integrated measures of viral diversity such as sequence composition, viral hosts, and phage lifestyles^9^, (see Figure 1) with social, pre-, peri- and postnatal environmental exposures. We revealed the effects of these exposures on viral community and the possible effects on metabolism.

In previous reports, *Crassvirales* (class *Caudoviricetes*) and *Microviridae* (class *Malgrandaviricetes*) phages were found to be the two most abundant viral groups in the adult human gut, with their relative abundance being negatively correlated^19,44–47^. Here, in one-year-old infants, a similar observation was made, members of the *Caudoviricetes* and *Malgrandaviricetes* classes were the most abundant phages.

Interestingly, ongoing exposures such as having older siblings and residential location, as well as past exposures (e.g., birth weight, preeclampsia) were linked with gut virome composition at one year of age. However, it is still possible that the prenatal and perinatal exposures still influenced the immune education earlier in life and remnants of the interplay are still tangible at 1-year of age^5^. Among the exposures significantly influencing the gut virome composition, the largest effect sizes were from residential location (rural vs. urban) and having older siblings (see Figure 2C and 2D). Interestingly, urbanization has been reported to have a significant impact on the composition of the adult viral community, with individuals living in urban areas having higher abundance of *Lactococcus* (family *Streptococcaceae*) phages^48^. The latter is presumably associated with the consumption of dairy products. We show that the living environment also affects the gut virome of infants, and that *Streptococcaceae* targeting phages are also more abundant in infants living in urban areas, possibly reflecting differences in dietary habits rather than residence *per se* (Figure 3).

Having older siblings influences the development of the bacterial community in early life^49–51^ and here we show that having older siblings is also associated with gut virome composition at one year of age. Importantly, from a translational angle, early-life exposures may affect the establishment of health phenotypes, such as the protective role of breastfeeding against eukaryotic-viral infections in the neonatal period^30^. Combining gut bacterial compositional data with gut virome composition (Figure S3A and S5B-C) in our cohort elucidates the co-abundance of phages and their hosts, underlying the role of phage-host interactions in shaping the GM. Most of these viruses (Figure S3A) have temperate lifestyles, as evidenced by the presence of genes coding for integrases. Thus, these temperate phages appear to have the ability to integrate their genome into the bacterial hosts and become prophages at some point.

Gut virome members have the potential to modulate biochemical processes^12,13,52^. The functional prediction of the genes derived from vOTUs co-varying with exposures, revealed up to 90% of genes with unknown functions. It emphasizes that proteins with yet uncharacterized functions are potentially playing a role in the regulation of human host phenotypes. Certain predicted gene functions linked to metabolic activities, such as alanine, aspartate and glutamate metabolism, amino sugar and nucleotide sugar metabolism and glycolysis/gluconeogenesis, which are likely associated with dietary intake and degradation of macronutrients, were associated with fish in the diet, birth weight, residence location and egg in the diet (Figure 4A). Maternal obesity alters fatty acid metabolism and changes in gene expression of lipid metabolism in infants, which cause a higher risk of developing obesity and its complications, neuropsychiatric disorders and asthma^53,54^. We find here that viral genes associated with normal weight mothers were predominantly enriched in fatty acid biosynthesis compared to obese mothers, which may be an intermediate pathway by which maternal obesity affects child health. In addition, for biotin metabolism, which is known to be impaired by severe obesity^55^, many phage genes are also observed to be enriched in infants from mothers with BMI below 25 in our data. The mothers enrolled in the cohort participated in a randomized clinical trial where they were randomized to receiving fish oil or a placebo from week 24 of pregnancy to one week after birth^38,56^. The design also examined the difference in vitamin D between high and standard doses^39^, which had no effect on the viral community in our analysis. Of note, the supplementation of fish oil during pregnancy was not found to influence the gut bacterial component at age one year. Here we report that the same intervention has some influence on the gut virome at age one year, but the effect is only borderline significant. The infants of mothers that received fish oil had viral genes involved in lysine biosynthesis, glycerophospholipid metabolism, and purine metabolism – metabolic activities that have been associated to fish oil supplementation^57,58^, but never attributed to gut virome composition. Interestingly, most of these metabolism-related genes were conserved across temperate vOTUs targeting *Ruminococcus*, *Faecalibacterium* and *Prevotella* spp. (Figure 4B). These genera have been consistently reported to be enriched in Danish and American subjects with a diet rich in carbohydrates, resistant starch, and fibers, and being determinants of the so-called *Prevotella*-enterotype^59,60^. The *Prevotella*-enterotype is established early in life (between 9-36 months of age)^61–63^ and have been previously suggested as markers of GM maturity at age one year^2,64,65^. Stokholm et al. (2018) reported delayed GM maturation as a risk factor for later development of asthma indicating the importance of these microbes for immune maturation.

Although our study is currently unable to assess how these gut virome associated genes are actively involved in either enhancing either phage or host fitness, or both, our data underlines the potential importance of bacteriophage-encoded metabolic genes and delivers an initial insight of the type of metabolic content conveyed by the gut virome in association to environmental variables.

In summary, our data provides detailed insight into the influence of common environmental factors that shape the gut virome during early life. We also uncover that key gut metabolic functions can be encoded by viral genes, which suggest that, in addition of shaping gut bacteriome composition, phages may directly play a role in metabolic activities.

## Methods

### Study participants and Ethics

Participants belong to the COPSAC2010 cohort^32^. Fecal samples for virome extraction sequencing and analysis were collected from all infants at age 1 year. The study was conducted in accordance with the Declaration of Helsinki and was approved by The National Committee on Health Research Ethics (H-B-2008-093) and the Danish Data Protection Agency (2015-41-3696). Both parents gave written informed consent before enrollment.

### Sample collection, sequencing, virome assembly

Preparation of fecal samples, and extraction and sequencing of virions was carried out using a previously described protocol^33^. Briefly, viral-associated DNA was subjected to short MDA amplification and libraries were prepared following using manufacturer’s procedures for the Nextera XT kit (FC-131-1096 Ilumina, California). Libraries were single-end high-throughput sequenced on the Illumina HiSeq X platform. Details of the pipeline for data processing, de-novo assembly, quality control, bacterial-host and lifestyle predictions, abundance-mapping (vOTU table), and taxonomy of complete and partial viral genomes (here termed vOTUs) can be found in Shah et al. (2021). 16S rRNA gene amplicon data (bOTU table) from the same cohort’s participants were retrieved from Stokholm et al. (2018).

### Environmental exposures

Briefly, during scheduled visits to the COPSAC clinic, information on a wide range of exposures was collected. A total of 30 environmental exposures were investigated and were grouped into social (n = 6), pre- (n = 4), peri- (n = 9) and postnatal (n = 11) exposures based on whether they occurred or existed before birth. See Supplementary Table S1 for a complete list of the exposures.

### Statistics and data analysis

Analyses on diversity were carried out on contingency tables gathering vOTUs abundance. Abundance data was normalized by reads per kilobase per million (RPKM). Alpha-diversity (Observed vOTUs and Shannon Index) indices and Beta-diversity (Bray-Curtis and Sørensen-Dice distances) matrices were generated using the package phyloseq (version 1.42.0)^66^. The contribution of each covariate to explain vOTUs community structure (as determined by Sørensen-Dice similarity and Bray-Curtis dissimilarity metrics) was calculated using distance-based redundancy analysis (db-RDA) models coupled to adonis PERMANOVA (n permutations = 999) in package vegan (version 2.6-2)^67^, while the effect size of the same covariates on alpha-diversity was calculated with linear mixed models from the package lmerTest (version 3.1-3)^68^. All linear mixed models accounted for technical variation between runs using sequencing lane as the random effect.

Different differential abundance analysis methods were evaluated by DAtest^69^. DESeq2 (version 1.36.0) performed well with a low false positive rate and a high ability to detect differential vOTUs for our data^70^. The sequencing lane was considered as a factor-covariate. The raw reads count table of each sample for vOTUs were prepared as input. All parameters are default except for sfType which is set to poscount. Benjamini and Hochberg method was adapted to correct the p-values. vOTUs with adjusted p-value ≤ 0.001 and log2 fold change ≥ |1| were selected for downstream analyses.

Spearman’s rank correlations were used to test univariate associations of continuous data, and results were visualized in a heatmap. MAFFT^71^ was used to generate the phylogenetic tree file for those highly correlated vOTUs. The phylogenetic tree was visualized using the R package ggtree (version 3.4.0)^72^. Procrustes analysis (R package vegan) was performed on vOTUs as target block and 16S rRNA gene data as rotatory block (n permutations = 999), while using the first two constrained components (CAP1 and CAP2) of db-RDA models for each data block.

ORF calling on selected vOTUs was executed with Prodigal^73^. To determine metabolic function, genes were annotated based on KEGG Orthology using KofamScan^41^ and filtered by default thresholds. Enricher function in clusterProfiler package (version 4.6.0) was applied to detect whether genes in differently abundant vOTUs were enriched in the metabolic pathway^74^.

All analyses were carried out in R (version 4.0.2) and results were visualized with the package ggplot2 (version 3.3.6)^75^.

## Supporting information

Supplemental results

## Data and code availability

Sequencing FASTQ files are available on ENA under project number PRJEB46943. All cohort participants’ individual-level data are protected by Danish and European law and are not publicly available. Codes for data analyses are available from the authors upon request.

## Acknowledgements

We express our deepest gratitude to the children and families of the COPSAC2010 cohort study for all their support and commitment. We acknowledge and appreciate the unique efforts of the COPSAC research team.

## Author contributions

S.M., M.A.P., J.S. and D.S.N. conceived the project and supervised all the research; B.C., K.B., J.S., S.J.S., L.J. and M.D. collected the samples and/or information; Y.Z., J.L.C.M., S.A.S. and D.S.N. analyzed the data; Y.Z., J.L.C.M. and D.S.N. wrote the manuscript with the assistance of L.D., S.A.S., J.T., C.L.R., S.M., M.A.P. and J.S.; L.D. prepared the virome and sequencing libraries; all authors contributed to, revised and approved the final manuscript.

## References

1. Tamburini, S., Shen, N., Wu, H. C. & Clemente, J. C. The microbiome in early life:implications for health outcomes. Nat. Med. 22, 713–722 (2016).

2. Stokholm, J. et al. Maturation of the gut microbiome and risk of asthma in childhood. Nat. Commun. 9, 141 (2018).

3. Pihl, A. F. et al. The role of the gut microbiota in childhood obesity. Child. Obes. 12, 292–299 (2016).

4. Kostic, A. D. et al. The dynamics of the human infant gut microbiome in development and in progression toward type 1 diabetes. Cell Host Microbe 17, 260–273 (2015).

5. Vatanen, T. et al. Variation in microbiome LPS immunogenicity contributes to autoimmunity in humans. Cell 165, 842–853 (2016).

6. McCann, A. et al. Viromes of one year old infants reveal the impact of birth mode on microbiome diversity. Peerj 6, e4694 (2018).

7. Sutton, T. D. S. & Hill, C. Gut bacteriophage: current understanding and challenges. Front. Endocrinol. 10, 784 (2019).

8. Shkoporov, A. N. & Hill, C. Bacteriophages of the human gut: the “known unknown’’ of the microbiome. Cell Host Microbe 25, 195–209 (2019).

9. Shah, S. A. et al. Expanding known viral diversity in the healthy infant gut. Nat. Microbiol., Accepted (2023).

10. Gregory, A. C. et al. The gut virome database reveals age-dependent patterns of virome diversity in the human gut. Cell Host Microbe 28, 724–740 (2020).

11. Roux, S. et al. IMG/VR v3: an integrated ecological and evolutionary framework for interrogating genomes of uncultivated viruses. Nucleic Acids Res. 49, D764–D775 (2021).

12. Kieft, K. et al. Ecology of inorganic sulfur auxiliary metabolism in widespread bacteriophages. Nat. Commun. 12, 3503 (2021).

13. Bi, L. et al. Diversity and potential biogeochemical impacts of viruses in bulk and rhizosphere soils. Environ. Microbiol. 23, 588–599 (2021).

14. Tisza, M. J. & Buck, C. B. A catalog of tens of thousands of viruses from human metagenomes reveals hidden associations with chronic diseases. P Natl Acad Sci USA 118, e2023202118 (2021).

15. Clooney, A. G. et al. Whole-virome analysis sheds light on viral dark matter in inflammatory bowel disease. Cell Host Microbe 26, 764–778 (2019).

16. Zhao, L. Y. et al. Uncovering 1,058 novel human enteric DNA viruses through deep long-read third-generation sequencing and their clinical impact. Gastroenterology 162, S96–S96 (2022).

17. Kaelin, E. A. et al. Longitudinal gut virome analysis identifies specific viral signatures that precede necrotizing enterocolitis onset in preterm infants. Nat. Microbiol. 7, 653–662 (2022).

18. Reyes, A. et al. Gut DNA viromes of Malawian twins discordant for severe acute malnutrition. P Natl Acad Sci USA 112, 11941–11946 (2015).

19. Zhao, G. Y. et al. Intestinal virome changes precede autoimmunity in type I diabetes-susceptible children. P Natl Acad Sci USA 114, E6166–E6175 (2017).

20. Vehik, K. et al. Prospective virome analyses in young children at increased genetic risk for type 1 diabetes. Nat. Med. 25, 1865–1872 (2019).

21. Tomofuji, Y. et al. Whole gut virome analysis of 476 Japanese revealed a link between phage and autoimmune disease. Ann. Rheum. Dis. 81, 278–288 (2021).

22. Rasmussen, T. S. et al. Faecal virome transplantation decreases symptoms of type 2 diabetes and obesity in a murine model. Gut 69, 2122–2130 (2020).

23. Ott, S. J. et al. Efficacy of sterile fecal filtrate transfer for treating patients with clostridium difficile infection. Gastroenterology 152, 799–811 (2017).

24. Brunse, A. et al. Fecal filtrate transplantation protects against necrotizing enterocolitis. ISME J 16, 686–694 (2022).

25. Stewart, C. J. et al. Temporal development of the gut microbiome in early childhood from the TEDDY study. Nature 562, 583–588 (2018).

26. Breitbart, M. et al. Viral diversity and dynamics in an infant gut. Res. Microbiol. 159, 367–373 (2008).

27. Lim, E. S. et al. Early life dynamics of the human gut virome and bacterial microbiome in infants. Nature Medicine 21, 1228–1234 (2015).

28. Taboada, B. et al. The gut virome of healthy children during the first year of life is diverse and dynamic. PLoS One 16, e0240958 (2021).

29. Liang, G. & Bushman, F. D. The human virome: assembly, composition and host interactions. Nat. Rev. Microbiol. 19, 514–527 (2021).

30. Liang, G. X. et al. The stepwise assembly of the neonatal virome is modulated by breastfeeding. Nature 581, 470–474 (2020).

31. Walters, W. A. et al. Longitudinal comparison of the developing gut virome in infants and their mothers. Cell Host Microbe 31, 187–198 (2023).

32. Bisgaard, H. et al. Deep phenotyping of the unselected COPSAC2010 birth cohort study. Clin. Exp. Allergy 43, 1384–1394 (2013).

33. Deng, L. et al. A protocol for extraction of infective viromes suitable for metagenomics sequencing from low volume fecal samples. Viruses 11, 667 (2019).

34. Edwards, R. A., McNair, K., Faust, K., Raes, J. & Dutilh, B. E. Computational approaches to predict bacteriophage-host relationships. FEMS Microbiol. Rev. 40, 258–272 (2016).

35. Pride, D. T., Wassenaar, T. M., Ghose, C. & Blaser, M. J. Evidence of host-virus co-evolution in tetranucleotide usage patterns of bacteriophages and eukaryotic viruses. BMC Genom. 7, 1–13 (2006).

36. Krupovic, M. & Forterre, P. Microviridae goes temperate: microvirus-related proviruses reside in the genomes of bacteroidetes. PLoS One 6, e19893 (2011).

37. Vinding, R. K. et al. Fish oil supplementation in pregnancy increases gestational age, size for gestational age, and birth weight in infants: a randomized controlled trial. J. Nutr. 149, 628–634 (2019).

38. Bisgaard, H. et al. Fish oil-derived fatty acids in pregnancy and wheeze and asthma in offspring. New Engl J Med 375, 2530–2539 (2016).

39. Chawes, B. L. et al. Effect of vitamin D-3 supplementation during pregnancy on risk of persistent wheeze in the offspring: a randomized clinical trial. JAMA 315, 353–361 (2016).

40. Li, X. J. et al. The infant gut resistome associates with E. coli, environmental exposures, gut microbiome maturity, and asthma-associated bacterial composition. Cell Host Microbe 29, 975–987 (2021).

41. Aramaki, T. et al. KofamKOALA: KEGG Ortholog assignment based on profile HMM and adaptive score threshold. Bioinformatics 36, 2251–2252 (2020).

42. Sanz-Gaitero, M., Seoane-Blanco, M. & van Raaij, M. J. Structure and function of bacteriophages. Bacteriophages: Biology, Technology, Therapy, 1–73 (2019).

43. Warwick-Dugdale, J., Buchholz, H. H., Allen, M. J. & Temperton, B. Host-hijacking and planktonic piracy: how phages command the microbial high seas. Virol. J. 16, 1–13 (2019).

44. Shkoporov, A. N. et al. The human gut virome is highly diverse, stable, and individual specific. Cell Host Microbe 26, 527–541 (2019).

45. Kim, K. H. & Bae, J. W. Amplification methods bias metagenomic libraries of uncultured single-stranded and double-stranded DNA viruses. Appl. Environ. Microbiol. 77, 7663–7668 (2011).

46. Roux, S. et al. Towards quantitative viromics for both double-stranded and single-stranded DNA viruses. Peerj 4, e2777 (2016).

47. Yutin, N. et al. Discovery of an expansive bacteriophage family that includes the most abundant viruses from the human gut. Nat Microbiol 3, 38–46 (2018).

48. Zuo, T. et al. Human-gut-DNA virome variations across geography, ethnicity, and urbanization. Cell Host Microbe 28, 741–751 (2020).

49. Laursen, M. F. et al. Having older siblings is associated with gut microbiota development during early childhood. BMC Microbiol. 15, 1–9 (2015).

50. Hedin, C. R., van der Gast, C. J., Stagg, A. J., Lindsay, J. O. & Whelan, K. The gut microbiota of siblings offers insights into microbial pathogenesis of inflammatory bowel disease. Gut Microbes 8, 359–365 (2017).

51. Christensen, E. D. et al. The developing airway and gut microbiota in early life is influenced by age of older siblings. Microbiome 10, 106 (2022).

52. Brown, E. M. et al. Gut microbiome ADP-ribosyltransferases are widespread phage-encoded fitness factors. Cell Host Microbe 29, 1351–1365 (2021).

53. Alvarez, D. et al. Impact of maternal obesity on the metabolism and bioavailability of polyunsaturated fatty acids during pregnancy and breastfeeding. Nutrients 13, 19 (2020).

54. Costa, S. M. et al. Maternal obesity programs mitochondrial and lipid metabolism gene expression in infant umbilical vein endothelial cells. Int J Obes (Lond) 40, 1627–1634 (2016).

55. Belda, E. et al. Impairment of gut microbial biotin metabolism and host biotin status in severe obesity: effect of biotin and prebiotic supplementation on improved metabolism. Gut 71, 2463–2480 (2022).

56. Hjelmso, M. H. et al. Prenatal dietary supplements influence the infant airway microbiota in a randomized factorial clinical trial. Nat. Commun. 11, 426 (2020).

57. Hishikawa, D., Valentine, W. J., Iizuka-Hishikawa, Y., Shindou, H. & Shimizu, T. Metabolism and functions of docosahexaenoic acid-containing membrane glycerophospholipids. FEBS Lett. 591, 2730–2744 (2017).

58. Cao, Y. et al. Integrated analysis of metabolomics and transcriptomics for assessing effects of fish meal and fish oil replacement on the metabolism of rainbow trout (Oncorhynchus mykiss). Front. Mar. Sci. 9, 208 (2022).

59. Roager, H. M. et al. Whole grain-rich diet reduces body weight and systemic low-grade inflammation without inducing major changes of the gut microbiome: a randomised cross-over trial. Gut 68, 83–93 (2019).

60. Wu, G. D. et al. Linking long-term dietary patterns with gut microbial enterotypes. Science 334, 105–108 (2011).

61. Bergstrom, A. et al. Establishment of intestinal microbiota during early life: a longitudinal, explorative study of a large cohort of danish infants. Appl. Environ. Microbiol. 80, 2889–2900 (2014).

62. Backhed, F. et al. Dynamics and stabilization of the human gut microbiome during the first year of life. Cell Host Microbe 17, 690–703 (2015).

63. Roswall, J. et al. Developmental trajectory of the healthy human gut microbiota during the first 5 years of life. Cell Host Microbe 29, 765–776 (2021).

64. Larsen, J. M. The immune response to Prevotella bacteria in chronic inflammatory disease. Immunology 151, 363–374 (2017).

65. Park, H., Shin, J. W., Park, S. G. & Kim, W. Microbial communities in the upper respiratory tract of patients with asthma and chronic obstructive pulmonary disease. PLoS One 9, e109710 (2014).

66. McMurdie, P. J. & Holmes, S. phyloseq: an R package for reproducible interactive analysis and graphics of microbiome census data. PLoS One 8, e61217 (2013).

67. Dixon, P. VEGAN, a package of R functions for community ecology. J. Veg. Sci. 14, 927–930 (2003).

68. Kuznetsova, A., Brockhoff, P. B. & Christensen, R. H. B. lmerTest Package: tests in linear mixed effects models. J. Stat. Softw. 82, 1–26 (2017).

69. Russel, J. et al. DAtest: a framework for choosing differential abundance or expression method. bioRxiv, 241802 (2018).

70. Love, M. I., Huber, W. & Anders, S. Moderated estimation of fold change and dispersion for RNA-seq data with DESeq2. Genome Biol. 15, 550 (2014).

71. Katoh, K. & Standley, D. M. MAFFT multiple sequence alignment software version 7: improvements in performance and usability. Mol. Biol. Evol. 30, 772–780 (2013).

72. Yu, G. C., Smith, D. K., Zhu, H. C., Guan, Y. & Lam, T. T. Y. GGTREE: an R package for visualization and annotation of phylogenetic trees with their covariates and other associated data. Methods Ecol. Evol. 8, 28–36 (2017).

73. Hyatt, D. et al. Prodigal: prokaryotic gene recognition and translation initiation site identification. BMC Bioinform. 11, 1–11 (2010).

74. Wu, T. et al. clusterProfiler 4.0: A universal enrichment tool for interpreting omics data. Innovation (Camb) 2, 100141 (2021).

75. Hadley, W. ggplot2: elegant graphics for data analysis. Springer-Verlag New York (2016).

