## Supplemental results for "The influence of early life exposures on the infant gut virome"

#### Supplementary Table 1

| Variable | Types | | | Sample numbers | | | Group |
|---|---|---|---|---|---|---|---|
| Education | Low | Medium | High | 52 | 412 | 181 | Social |
| Income | Normal | High | | 404 | 240 | | Social |
| Maternal age (median 32.07) | Below median | Above median | | 323 | 322 | | Social |
| Maternal BMI | < 25 | ≥ 25 | | 410 | 231 | | Social |
| Race | Caucasian | Non-caucasian | | 615 | 30 | | Social |
| AB usage during pregnancy | No | Yes | | 410 | 234 | | Prenatal |
| Fishoil during pregnancy | No | Yes | | 324 | 320 | | Prenatal |
| Preeclampsia | No | Yes | | 615 | 30 | | Prenatal |
| Smoking during pregnancy | No | Yes | | 599 | 46 | | Prenatal |
| AB usage during birth | No | Yes | | 627 | 18 | | Perinatal |
| Birth induction | No | Yes | | 426 | 219 | | Perinatal |
| Delivery method | Vaginal | CS | | 509 | 136 | | Perinatal |
| Gestational age (weeks) | < 37 | ≥ 37 | | 25 | 620 | | Perinatal |
| Hospitalized after birth | No | Yes | | 571 | 74 | | Perinatal |
| Low birth weight (< 2.5kg) | No | Yes | | 626 | 19 | | Perinatal |
| Mother AB usage during birth | No | Yes | | 440 | 202 | | Perinatal |
| Sex | Female | Male | | 310 | 335 | | Perinatal |
| Weight at birth | < 3.5 | ≥ 3.5 | | 285 | 360 | | Perinatal |
| AB usage before 6 months old | No | Yes | | 560 | 79 | | Postnatal |
| AB usage before 1 year old | No | Yes | | 343 | 296 | | Postnatal |
| Cat or dog at home | No | Yes | | 420 | 222 | | Postnatal |
| Cow milk introduced in diet, age in days (median 214 days) | Below median | Above median | | 325 | 320 | | Postnatal |
| Daycare at sampling time | No | Yes | | 132 | 511 | | Postnatal |
| Egg introduced in diet, age in days (median 257 days) | Below median | Above median | | 323 | 322 | | Postnatal |
| Fish introduced in diet, age in days (median 212 days) | Below median | Above median | | 325 | 320 | | Postnatal |
| Older siblings | No | Yes | | 286 | 359 | | Postnatal |
| Duration of exclusion breastfeeding | < 120 | ≥ 120 | | 277 | 368 | | Postnatal |
| Still breast fed at sampling time | Mixed | No breastfeeding | | 93 | 550 | | Postnatal |
| Type of living environment | Urban | Rural | | 304 | 340 | | Postnatal |

**Supplementary Table S1**

Environmental exposure characteristics of the infant samples. Column "types" shows the composition of each exposure. Column "numbers" depicts the number of samples (N) for each factor. The group column shows how each variable is grouped.

### Supplementary Table 2

| | Host phylum | Host family | Number of vOTUs | | | Virulent/ temperate |
|---|---|---|---|---|---|---|
| | | | temperate | virulent | unknown | |
| 1 | *Actinobacteria* | *Bifidobacteriaceae* | 253 | 80 | 221 | 0.32 |
| 2 | *Actinobacteria* | Other | 33 | 58 | 58 | 1.76 |
| 3 | *Bacteroidetes* | *Bacteroidaceae* | 238 | 180 | 1035 | 0.76 |
| 4 | *Bacteroidetes* | *Barnesiellaceae* | 6 | 1 | 39 | 0.17 |
| 5 | *Bacteroidetes* | *Flavobacteriaceae* | 4 | 11 | 38 | 2.75 |
| 6 | *Bacteroidetes* | *Odoribacteraceae* | 2 | 7 | 25 | 3.50 |
| 7 | *Bacteroidetes* | *Prevotellaceae* | 42 | 15 | 148 | 0.36 |
| 8 | *Bacteroidetes* | *Rikenellaceae* | 11 | 8 | 58 | 0.73 |
| 9 | *Bacteroidetes* | *Tannerellaceae* | 41 | 66 | 186 | 1.61 |
| 10 | *Bacteroidetes* | Other | 20 | 89 | 140 | 4.45 |
| 12 | *Firmicutes* | *Clostridiaceae* | 257 | 49 | 332 | 0.19 |
| 13 | *Firmicutes* | *Erysipelotrichaceae* | 101 | 19 | 133 | 0.19 |
| 14 | *Firmicutes* | *Lachnospiraceae* | 1058 | 63 | 1066 | 0.06 |
| 15 | *Firmicutes* | *Paenibacillaceae* | 47 | 1 | 138 | 0.02 |
| 16 | *Firmicutes* | *Peptostreptococcaceae* | 99 | 2 | 182 | 0.02 |
| 17 | *Firmicutes* | *Ruminococcaceae* | 672 | 13 | 1089 | 0.02 |
| 18 | *Firmicutes* | *Streptococcaceae* | 48 | 26 | 181 | 0.54 |
| 19 | *Firmicutes* | *Veillonellaceae* | 399 | 0 | 303 | 0.00 |
| 20 | *Firmicutes* | Other | 778 | 32 | 1236 | 0.04 |
| 21 | *Proteobacteria* | *Enterobacteriaceae* | 160 | 0 | 316 | 0.00 |
| 22 | *Proteobacteria* | *Sutterellaceae* | 59 | 0 | 111 | 0.00 |
| 23 | *Proteobacteria* | Other | 78 | 38 | 285 | 0.49 |
| 24 | *Verrucomicrobia* | *Akkermansiaceae* | 67 | 17 | 72 | 0.25 |
| 26 | Other | Other | 466 | 220 | 1815 | 0.47 |

**Supplementary Table S2**

Number of different lifestyle vOTUs.

#### Supplementary Figure 1

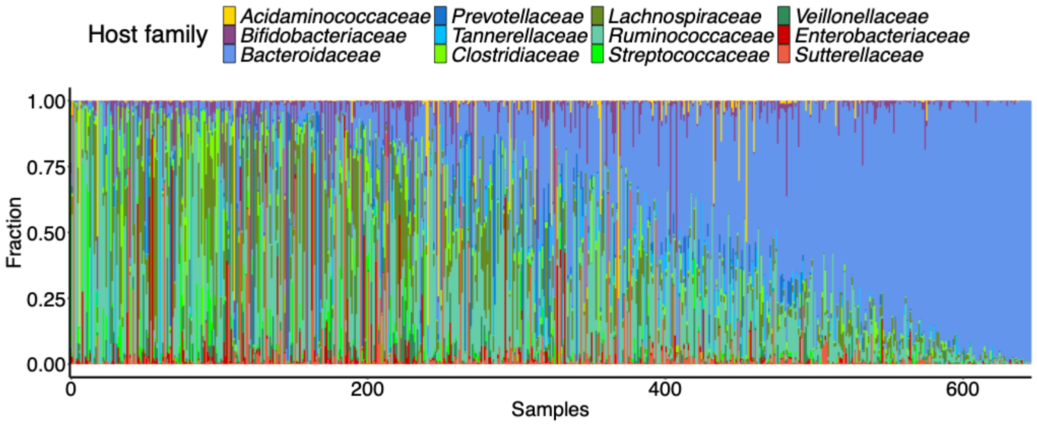


**Supplementary Figure S1**

Relative abundance of vOTUs across all samples grouped by their host bacterial family.

#### Supplementary Figure 2

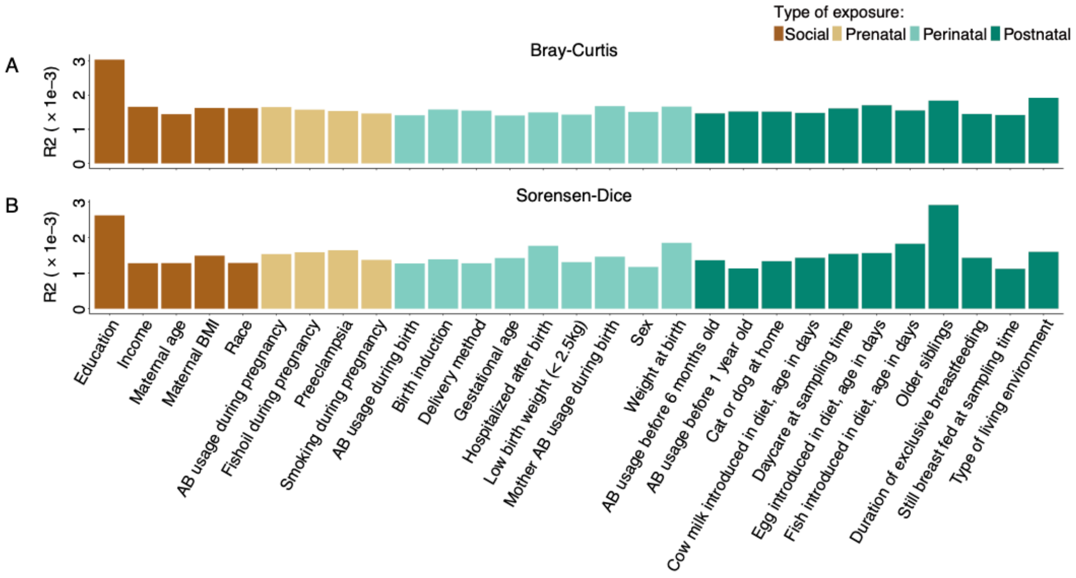


**Supplementary Figure S2**

A-B) The effect size of exposures on virome variation calculated by db-RDA on Bray-Curtis (A) and Sorensen-Dice (B) dissimilarity matrices.

Supplementary Figure 3

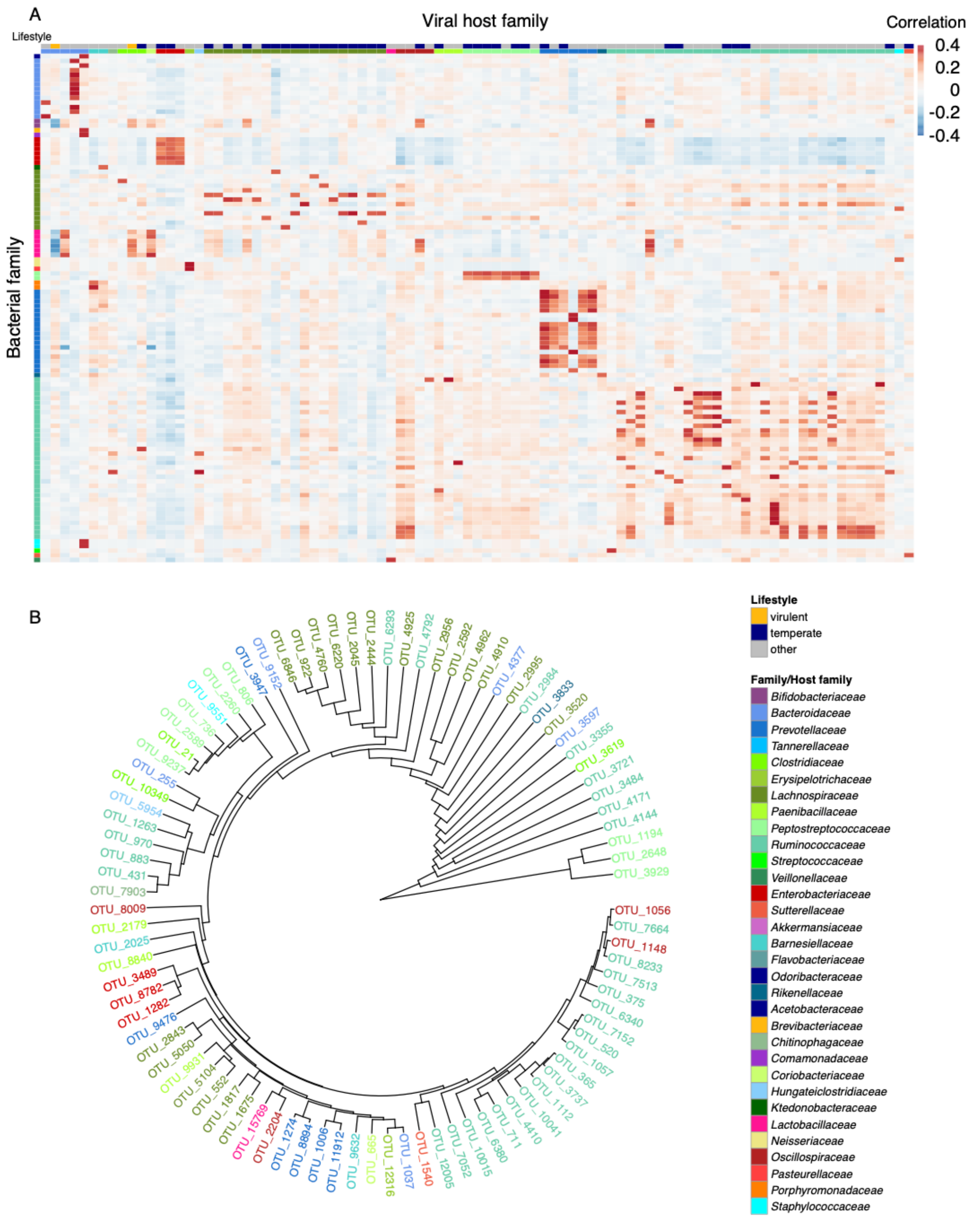


**Supplementary Figure 3**

A) Heatmap of Spearman's rank correlation between the differentially abundant vOTUs and 16S rRNA (V4 region) data blocks. The rows represent the 91 vOTUs, and the columns represent the 110 bacterial OTUs. Virome contigs were labeled according to their host family. Bacterial OTUs were labeled according to their family.

B) Phylogenetic tree showing genetic relationships of vOTUs.

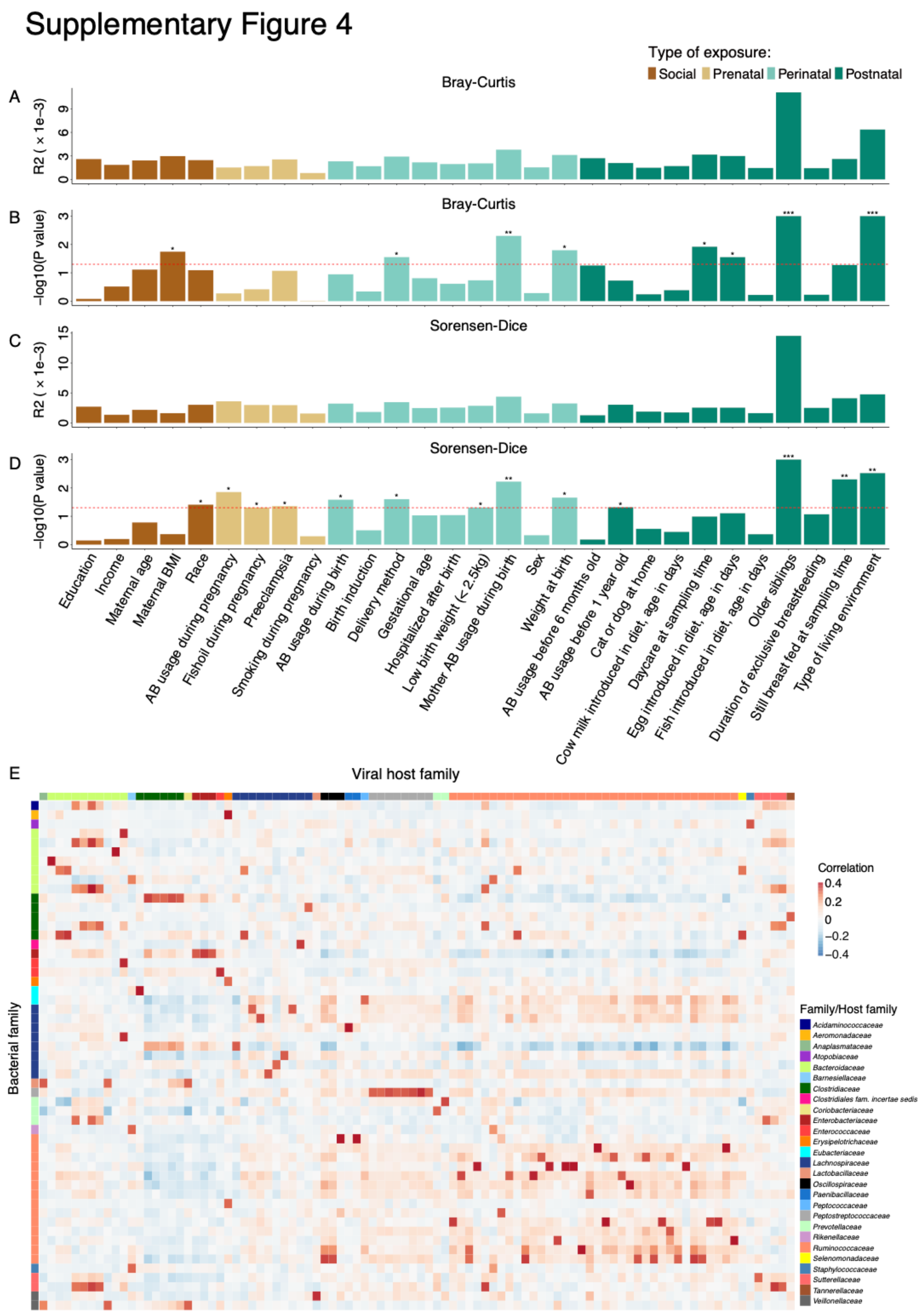


**Supplementary Figure 4**

A-D) Barplots showing p-values and effect sizes of db-RDA analysis for bacterial variation based on Bray-Curtis (A-B) and Sorensen-Dice (C-D) dissimilarity matrices. P-values were calculated by an ANOVA-like permutation test (n = 999).

E) Heatmap of Spearman's rank correlation between the differentially abundant vOTUs and whole-genome shotgun metagenome data blocks.

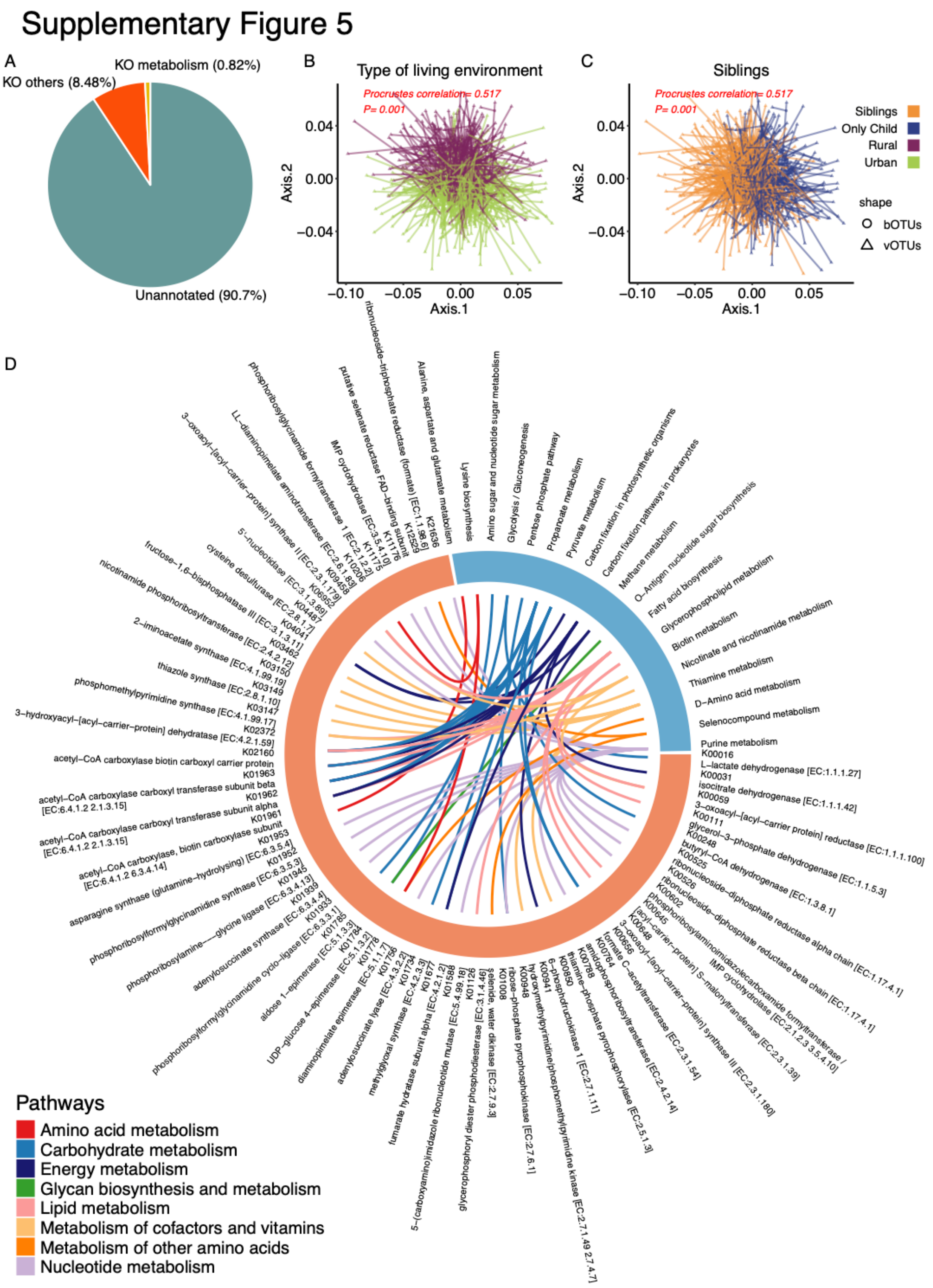


**Supplementary Figure 5**

A) Distribution of gene annotations to KEGG databases.

B-C) Procrustes correlation between the virome data and bacterial 16S rRNA gene data. Sorensen-Dice was used to generate distance matrices, the results of db-RDA were taken into the procrustes analysis. The triangle represents the virome samples and the circle represents the bacteria samples.

D) KOs (enzymes and E.C. numbers) that are associated with metabolism pathways.
